# Formation of the β-sheet at the extracellular opening of the intimin β-barrel domain is necessary for stability and efficient passenger secretion

**DOI:** 10.1101/2025.05.11.653311

**Authors:** Sai Priya Sarma Kandanur, Lara Breuer, Fabian A. Renschler, Monika S. Schütz, Jack C. Leo

## Abstract

Attaching and effacing pathogens, such as enteropathogenic and enterohaemorrhagic *E. coli*, rely on an adhesin, intimin, for attachment to enterocytes and downstream effects leading to actin pedestal formation. Intimin belongs to the inverse autotransporter family (type 5e secretion systems), which secrete the extracellular adhesive domain or passenger via a hairpin intermediate. While many of the next steps in secretion are now understood, how the hairpin initially forms is not known. We sought to investigate this by making point mutations at several positions in the β-barrel domain of intimin, as this domain forms at least part of the secretion pore. We made the mutations in a wild-type background and in a stalled intermediate caught in the hairpin conformation, which allowed us to uncouple passenger secretion from hairpin formation. Surprisingly, most of the point mutations did not have an appreciable effect on hairpin formation or passenger secretion, and larger changes such as replacing the entire linker region with a flexible glycine-serine stretch showed only a modest reduction in passenger secretion. By contrast, mutations affecting a small β-sheet at the extracellular face of the intimin β-barrel between two extracellular loops and the C-terminus of the linker had a more pronounced effect on secretion, and abolishing this β-sheet prevented hairpin formation and led to complete loss of the wild-type protein. Our results show that the intimin β-barrel is remarkably tolerant to changes and that the β-sheet on the extracellular side of this domain plays a central role in passenger secretion and protein stability.

## Introduction

Intimin is a virulence factor of attaching and effacing pathogens, which include enteropathogenic and enterohaemorrhagic *Escherichia coli* (EPEC and EHEC, respectively). Intimin mediates the adhesion of these pathogens to enterocytes, which leads to a signal transduction cascade resulting in cytoskeletal rearrangements, the loss of microvilli and the formation of an actin pedestal around the bacteria [1]. Remarkably, the receptor for intimin is not a host protein but the translocated intimin receptor (Tir), which is translocated into host cells via a type 3 secretion system [2].

Intimin itself is an outer membrane protein with an extended extracellular region at the C-terminus including the Tir-binding domain [3], a transmembrane β-barrel domain [4], and a short periplasmic region contain a peptidoglycan-binding LysM domain [5]. The extracellular region or passenger consists of four immunoglobulin (Ig)-like domains and C-type lectin-like domain at the very C-terminus [3,6]. The passenger is connected to the β-barrel by a linker region that traverses the lumen of the β-barrel [4].

Over the past decade, it has become apparent that intimin is the prototype of a subclass of autotransporter proteins, termed type 5e secretion systems [7,8]. These are also referred to as inverse autotransporters because the domain order (N-terminal β-barrel and C-terminal passenger) is reversed when compared with classical (type 5a) autotransporters, which have an N-terminal passenger and a C-terminal β-barrel [9]. The mechanism of autotransport begins with the insertion of the intimin β-barrel into the outer membrane by the β-barrel assembly machinery (BAM) [7]. Probably concomitantly, a hairpin structure is formed between the linker and the N-terminal part of the passenger, positioning the very N-terminus of the passenger at the cell surface [10]. The hairpin occupies the pore of the β-barrel, or more probably a hybrid pore formed by intimin and the central component of the BAM, BamA, as has been demonstrated for classical autotransporters [11,12]. The hairpin allows the first extracellular Ig-like domain D00 to fold at the extracellular surface, pulling more of the passenger through the pore until the next Ig-like domain D0 can fold, which in turn pulls through the next domain. Thus, the secretion is driven by sequential folding of Ig-like domains and the free energy for translocating the polypeptide through the pore is provided by the energy released when the domains in the passenger fold [13].

Although the mechanism of passenger secretion is now well understood, it is not known how the hairpin is formed in the first place. To address this issue, we undertook a mutagenesis study to try to pinpoint residues in the β-barrel domain that would be required for efficient hairpin formation and passenger secretion. To this end, we employed a stalled variant of intimin, IntHA453, where a double haemagglutinin (HA) tag was inserted into the D00 domain [10]. This prevents the folding of the D00 domain and traps the passenger at the hairpin stage [13]. Using this stalled variant allowed us to uncouple hairpin formation from passenger secretion The HA tag can still be detected at the cell surface in IntHA453, demonstrating the formation of the hairpin. We introduced a number of point mutations or larger-scale changes to conserved features in the β-barrel domain in both the wild-type (WT) intimin and IntHA453 backgrounds and examined surface exposure of either the C-terminus of the passenger (in the WT background) or the double HA tag as a proxy for hairpin formation in the IntHA453 background. We found that hairpin formation and passenger secretion are remarkably robust, and most alterations had little to no effect. However, changes to the small β-sheet at the extracellular surface of the β-barrel which forms between the linker and two extracellular loops caused secretion defects or loss of the entire protein. Our results suggest this β-sheet is crucial for intimin stability and secretion.

## Results

### Identification of interactions in the intimin β-barrel domain

To investigate hairpin formation, we decided to target interactions between residues in the linker and residues lining the lumen of the β-barrel or in the surface loops. These were based on the existing crystal structure of the intimin β-barrel [4] We reasoned that such interactions would be needed to stabilise the linker and may contribute to hairpin formation. We focussed on three regions of the β-barrel: the periplasmic face, with an *α*-helical turn that plugs the bottom of the β-barrel, the linker itself as it traverses the pore of the β-barrel, and the small antiparallel β-sheet formed between two extracellular loops (loops 4 and 5) and the linker as it exits the pore (Figure 1). We made changes to these regions by site-directed mutagenesis, targeting conserved residues (Figure 1B).

**Figure 1.**
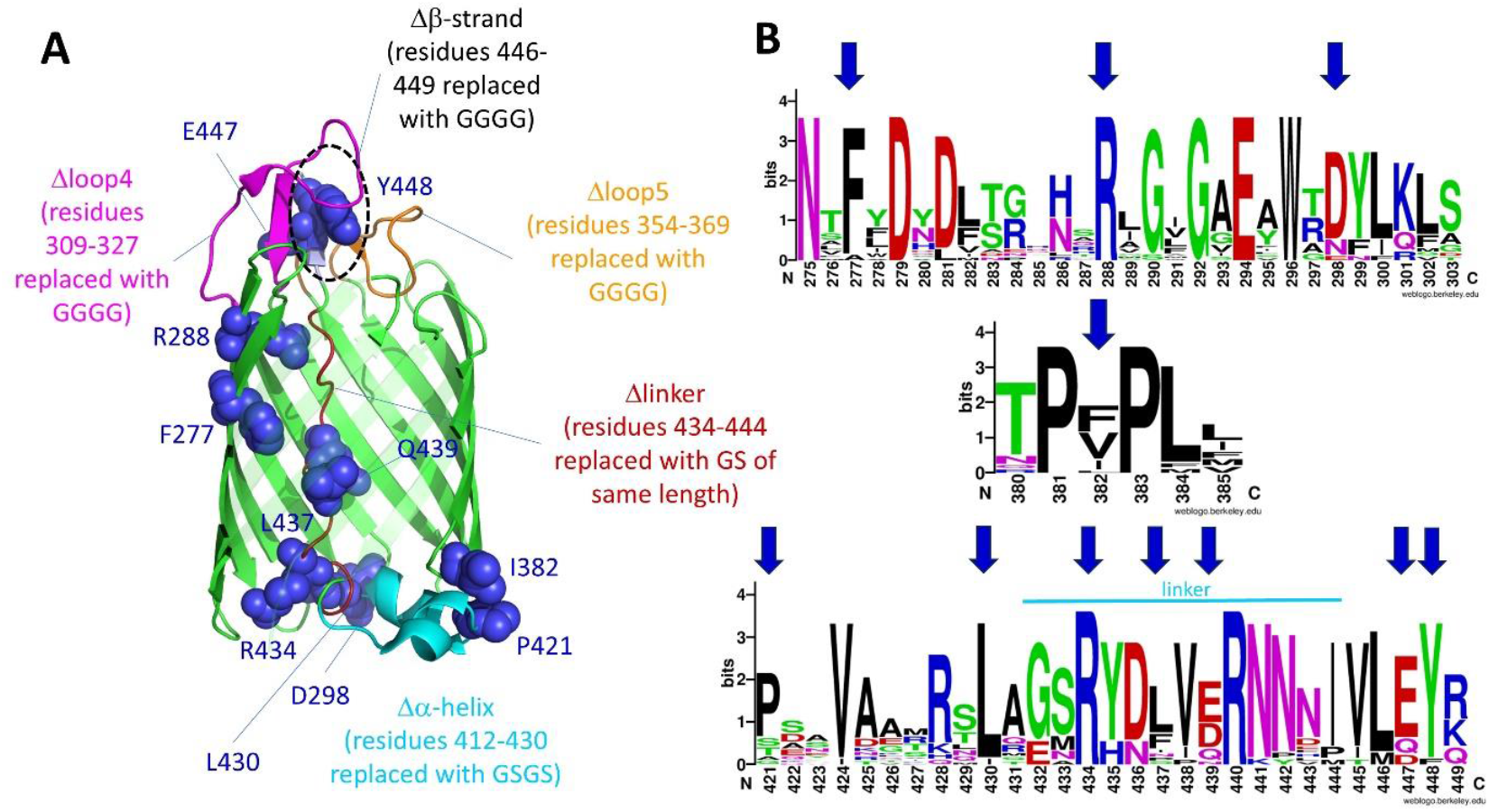
Residues in the intimin β-barrel domain targeted for mutation. A) The residues targeted for point mutation in the intimin β-barrel are highlighted as blue spheres. Larger mutations are highlighted in different colours (Δ*α*-helix, cyan; Δlinker, dark red; Δloop4, magenta; Δloop5, orange) and the Δβ-strand is encircled in black. Detailed interactions of the targeted residues and regions are given in Supplementary Figures 1-3. B) Conservation of the targeted residues in inverse autotranporters. Sequence logos based on an alignment of 20 inverse autotransporter sequences. Targeted residues are highlighted with arrows.

In the first region, we targeted residues D298, I382, P421, and L430. D298, located on periplasmic turn 3 of the β-barrel and facing the lumen, makes a hydrogen bond with the amide nitrogen of L430, potentially stabilising the N-terminus of the linker (Supplementary Figure 1). Changing this residue to alanine would disrupt this interaction. I382, at the C-terminal end of β-strand 14, forms the core of a hydrophobic cluster involving residues L384, V385, I419 and P421 (Supplementary Figure 1). The latter two are located on the *α*-helical turn. We reasoned that disrupting this hydrophobic cluster by changing I382 to a charged residue (aspartate) would destabilise the *α*-helical turn. P421 is located between the two short *α*-helices in the turn (Supplementary Figure 1). As a known secondary structure breaker, the proline at this position may be important to introduce the kink between the *α*-helices, so changing it to alanine may have an impact on the structure of this region. L430 is involved in a hydrophobic interaction with F266 on β-strand 4 (Supplementary Figure 1). This interaction might be important in positioning the N-terminus of the linker, and changing L430 to alanine should disrupt this interaction. In addition to these point mutations, we changed the entire *α*-helical turn (referred to as Δ*α*-helix, residues 412-430) with a flexible (serine-glycine)_2_ stretch, which is shorter than the *α*-helical turn but long enough to allow the linker to form in the correct position.

In the linker, we changed residues F277, R288, R434, L437 and Q439, all to alanine. F277 is located on β-strand 4 and packs against V438, located in the linker (Supplementary Figure 2); substituting F277 with alanine would abrogate this interaction. R288, on β-strand 6, is part of a hydrogen bonding network involving N441 and N443 at the C-terminus of the linker and Y353 on β-strand 9 (Supplementary Figure 2). This network would be lost upon exchanging R288 (completely conserved in inverse autotransporters, Figure 1B) to alanine. R434, located in the linker, is also completely conserved and interacts with the amide of E238 in intracellular turn 1 and has a cation-pi interaction with F266 on β-strand 4 (Supplementary Figure 1). L437 packs between L231 on β-strand 2 and the β-methylene groups of E214 and D229, on β-strands 1 and 2 respectively (Supplementary Figure 2); a substitution of L437 with alanine would disrupt these interactions. Q439 forms hydrogen bonds with K301 and the carbonyl group of S303 on β-strand 7. As for the *α*-helical turn, we assessed the global role of the linker by replacing all residues (434-444) with a glycine-serine stretch of the same length (referred to as Δlinker).

At the extracellular face of the β-barrel, we replaced E447 with alanine and Y448 with aspartate. Both of these residues are in the β-strand at the C-terminal tip of the linker. E447 makes a salt bridge with R325 and contacts R309, N328, N357, and Y354 through a bridging water molecule (Supplementary Figure 3). The highly conserved Y448 (Figure 1B) is at the centre of a hydrophobic cluster along with K319, Y322, A359, S363, and L446 that is expected to be disrupted in the Y448D mutation. In addition, Y448 forms a hydrogen bond to N318 on loop 4 (Supplementary Figure 3). We also changed this short β-strand (residues 446-449) entirely into glycines (Δβ-strand). In addition, we deleted extracellular loop 4 (Δloop4) by replacing residues 309-369 with four glycines, and loop 5 (Δloop5) by replacing residues 354-369 by four glycines. These four remaining, flexible residues would be able to connect to the adjacent β-strand but this results in fully deleting the extracellular loops.

### Mutations at the periplasmic face of the intimin β-barrel do not affect hairpin formation or passenger secretion

We introduced these mutations into both WT intimin (Int-Strep) and Intimin HA453 (IntHA453-Strep), both including a C-terminal StrepII tag. We tested the surface exposure of the passenger by immunofluorescence microscopy using an antibody against the C-terminus of intimin: as expected, cells producing Int-Strep were stained, whereas cells producing IntHA453-Strep did not stain with this antibody (Figure 2A). However, an anti-HA antibody did stain IntHA453-Strep, demonstrating that the double HA tag is surface-exposed and the hairpin is formed, as shown before [10]. A western blot with an anti-Strep tag antibody demonstrates that the Int-HA453-HA protein is intact, though expressed at lower levels than WT Int-Strep (Figure 2B). Both proteins exhibit the characteristic gel shift upon boiling that indicates unfolding of the β-barrel, which shows both proteins are correctly folded and inserted into the outer membrane. The blot also shows an additional, higher molecular-weight band (indicated by an asterisk in Figure 2B), which has been seen before [8,10,13] but the nature of which remains elusive. It is, however, clear that this band is not due to incorrect disulphide formation in the C-terminal domain of intimin, as addition of a reducing agent had no effect on this band (Supplementary Figure 4). Similarly, probing with the anti-Int antibody shows that IntHA453-Strep is produced at lower levels but that it is intact (Supplementary Figure 5). The lower expression of IntHA453-Strep seen in this study is possibly due to the autoinduction conditions used. With a longer expression time, the association of IntHA453 with the BAM complex [10] might lead to a reduction in outer membrane insertion efficiency over time. The empty vector control (pET22) failed to stain with either the anti-Intimin or the anti-HA antibody (Figure 2A).

**Figure 2.**
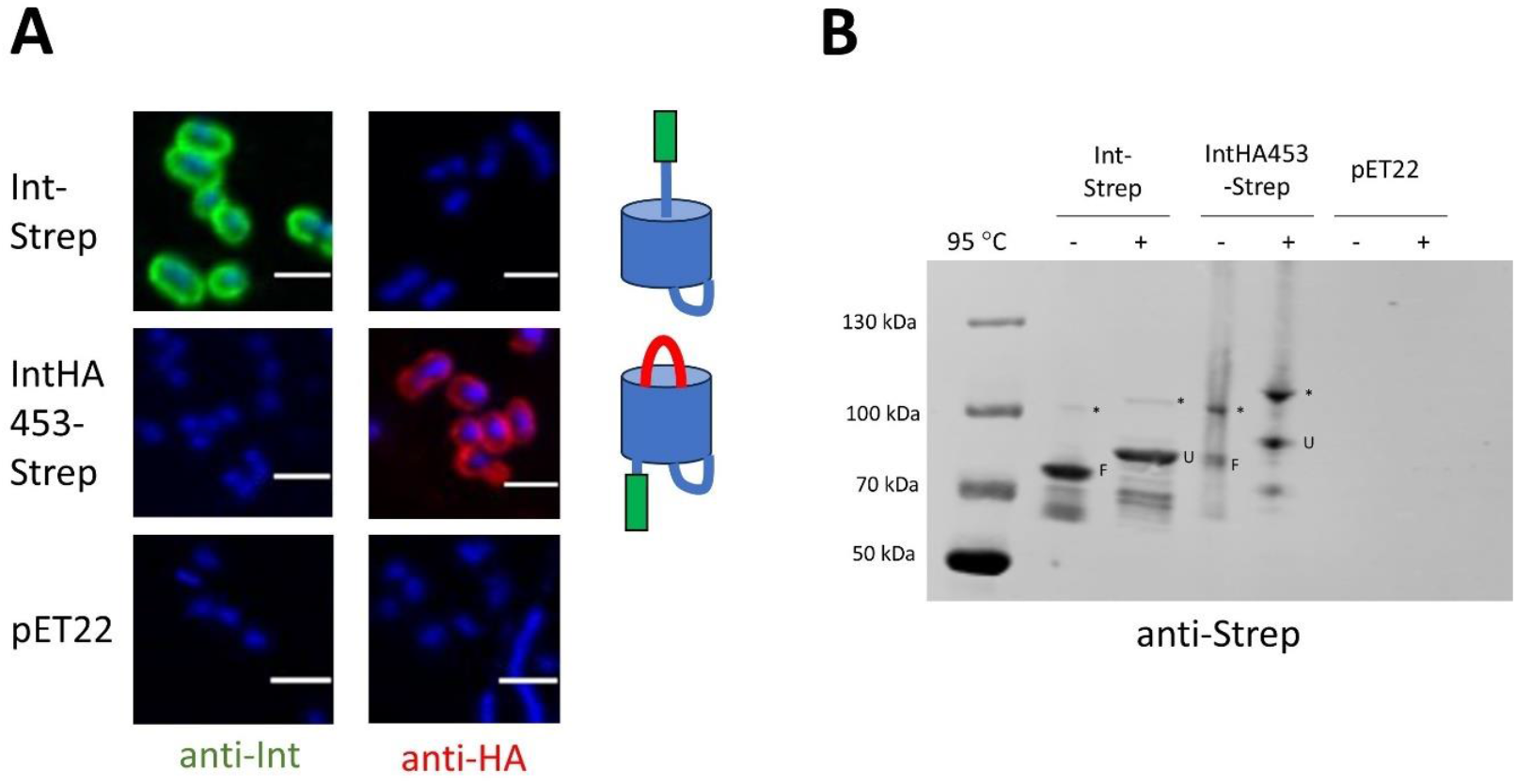
Surface exposure of intimin variants in autoinduced cultures. A) Confocal microscopy of cultures stained with an anti-intimin (anti-Int, left) or anti-HA tag (anti-HA, right) antibody. Cells were counterstained with DAPI. The schematics on the right of the microscopy images show the expected conformation of the proteins. pET22 is the vector control. The scale bar is 2 µm. B) Western blot of outer membrane samples with intimin variants using an anti-Strep tag primary antibody. Samples were split in half and incubated at either room temperature or 95 °C for 10 minutes and run in a 4-12% Novex gel to display the heat shift typical of β-barrel proteins before transferring to a nitrocellulose membrane. Molecular weight standards are notated on the left. F = folded, U= unfolded, * = band of unknown origin.

We then proceeded to examine the surface exposure of the various mutants, beginning with those on the periplasmic face of intimin. Remarkably, none of the mutations had a large effect on surface exposure of either the passenger in the Int-Strep background, or of the double HA tag in the IntHA453-Strep background (Figure 3A). All the proteins were produced, as verified by western blot, and all the mutations in the Int-Strep background exhibited a heat shift (Figure 3B), which suggests all the proteins were correctly folded and inserted into the outer membrane [14]. Interestingly, upon heating, some mutants (I382D, L430A) seemed to preferentially populate the higher molecular-weight band (denoted by an asterisk in Figures 3-5). Similarly to the WT IntHA453-Strep, mutants in this background were produced at lower levels than Int-Strep (Figure 3B). An exception to this seemed to be the Δ*α*-helix mutant, but this (like the I382D mutant) appeared to preferentially populate the higher molecular weight bands in the gel. Nonetheless, the mutant proteins all exhibited a heat shift (Figure 3B) and they were produced at high enough levels to be easily detected with the anti-HA antibody in immunofluorescence microscopy (Figure 3A). These results suggest that interactions at the periplasmic face of the intimin β-barrel are not important for efficient hairpin formation and passenger secretion.

**Figure 3.**
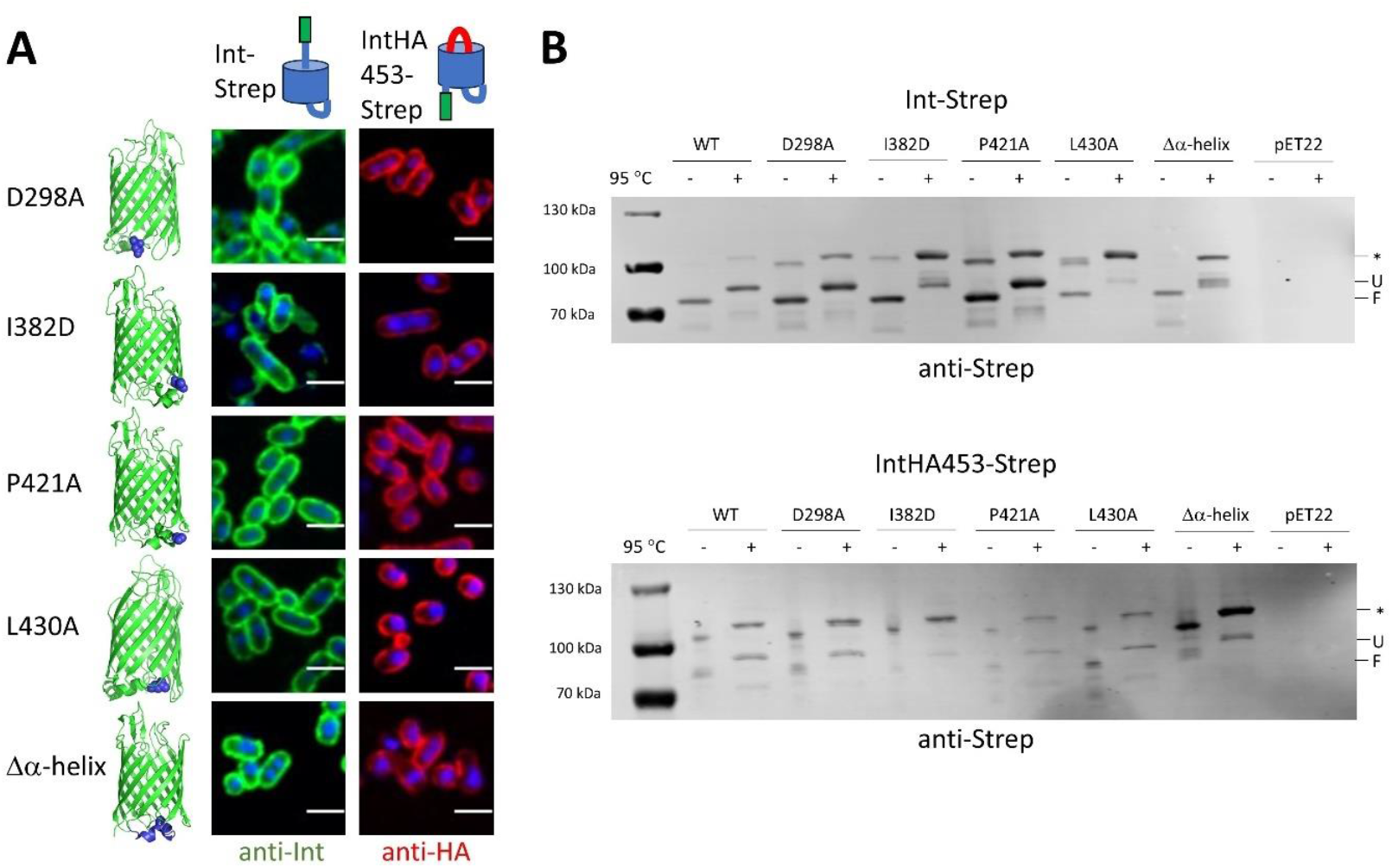
Effect of mutations on the periplasmic face of intimin. A) Confocal microscopy of intimin mutants in both the Int-Strep (left, probed with an anti-Strep antibody, coloured in green) and IntHA453-Strep (right, probed with an anti-HA antibody, coloured in red) backgrounds. Cells were counterstained with DAPI. Structures on the left highlight the position of the mutation. Schematics at the top show the expected conformation of the proteins. B) Western blots of outer membrane samples with intimin variants using an anti-Strep tag primary antibody. The upper blot is in the Int-Strep background and the lower blot in the IntHA453-Strep background. Samples were split in half and incubated at either room temperature or 95 °C for 10 minutes and run in a 4-12% Novex gel to display the heat shift typical of β-barrel proteins before transferring to a nitrocellulose membrane. pET22 is the empty vector. Molecular weight standards are notated on the left of the gels. F = folded, U= unfolded, * = band of unknown origin.

### Substituting the entire linker reduces secretion and hairpin formation

We then investigated whether interactions between the linker and the luminal face of the β-barrel affect hairpin formation and secretion. Similarly to our findings regarding the *α*-helical turn, individual point mutations in this region did not have an impact on passenger secretion or hairpin formation (Figure 4A). Only when the entire linker was replaced by a flexible sequence made of GS repeats did we observe an effect. Surface exposure of the passenger was reduced in the Int-Strep Δlinker variant in both backgrounds. Consistent with this observation, the amount of protein was reduced for Int-Strep Δlinker in the western blot, though the heat shift was still evident (Fig. 4B, upper panel). The reduction was also not due to degradation by the periplasmic protease DegP, which is known to play a role in inverse autotransporter quality control [7,15-17] (Supplementary Figure 6). All the other Int-Strep mutants behaved like the WT in the western blot (Figure 4B). Similarly to the previous set of mutants in the IntHA453, protein levels were reduced compared to WT Int-Strep, but all mutants displayed a heat shift (Fig. 4B, lower panel).

**Figure 4.**
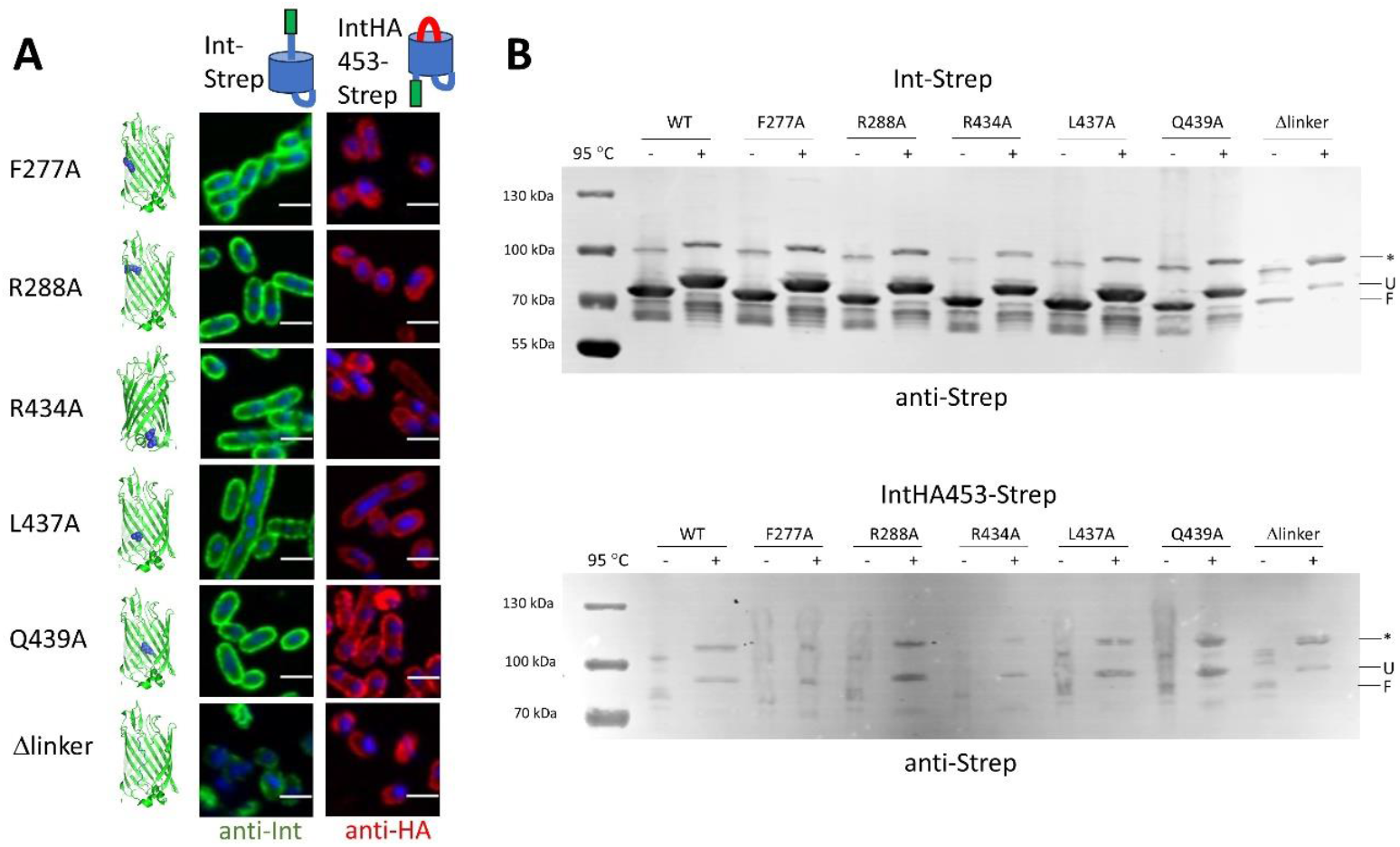
Effect of mutations on the linker region of intimin. A) Confocal microscopy of intimin mutants in both the Int-Strep (left, probed with an anti-Strep antibody, coloured in green) and IntHA453-Strep (right, probed with an anti-HA antibody, coloured in red) backgrounds. Cells were counterstained with DAPI. Structures on the left highlight the position of the mutation. Schematics at the top show the expected conformation of the proteins. B) Western blots of outer membrane samples with intimin variants using an anti-Strep tag primary antibody. The upper blot is in the Int-Strep background and the lower blot in the IntHA453-Strep background. Samples were split in half and incubated at either room temperature or 95 °C for 10 minutes to display the heat shift typical of β-barrel proteins and run in a 4-12% Novex gel to display the heat shift typical of β-barrel proteins before transferring to a nitrocellulose membrane. Molecular weight standards are notated on the left of the gels. F = folded, U= unfolded, * = band of unknown origin.

### Changes to the β-sheet at the extracellular face of the β-barrel affect secretion intimin stability

We examined the effect of changes in the β-sheet at the extracellular side of the β-barrel by making changes to residues involved in interactions presumed to stabilise the β-sheet. Though the E447A mutation did not have an effect on either hairpin formation or passenger secretion, the Y448D mutation did have a clear effect on hairpin formation, and the IntHA453-Strep Y448D-expressing cells failed to stain with the HA-antibody (Figure 5A). This was despite the protein being produced and exhibiting a normal heat shift (Figure 5B). Surprisingly, replacing the extracellular loops 4 and 5 with a shorter stretch of glycines did not have a large effect on either passenger secretion or hairpin formation (Figure 5A), though removing the longer loop 4 did lead to an increase in the higher molecular-weight band in the western blot (Figure 5B). By contrast, replacing the β-strand at the C-terminus of the linker with glycines had a dramatic effect, completely abolishing both hairpin formation and passenger secretion (Figure 5A). This was due to no protein being present in the outer membrane for the Int-Strep Δβ-strand mutant (Figure 5B), which we interpret to mean that the protein was destabilised and completely degraded. No protein was detected in whole cell extracts either, even when the periplasmic protease DegP was absent, demonstrating the importance of this β-strand for protein stability (Supplementary Figure 6). However, in the IntHA453-Strep variant, neither the Y448D nor the Δβ-strand mutation resulted in loss of the protein, though the HA tag could not be detected at the cell surface, suggesting the hairpin was not correctly formed (Figure 5B).

**Figure 5.**
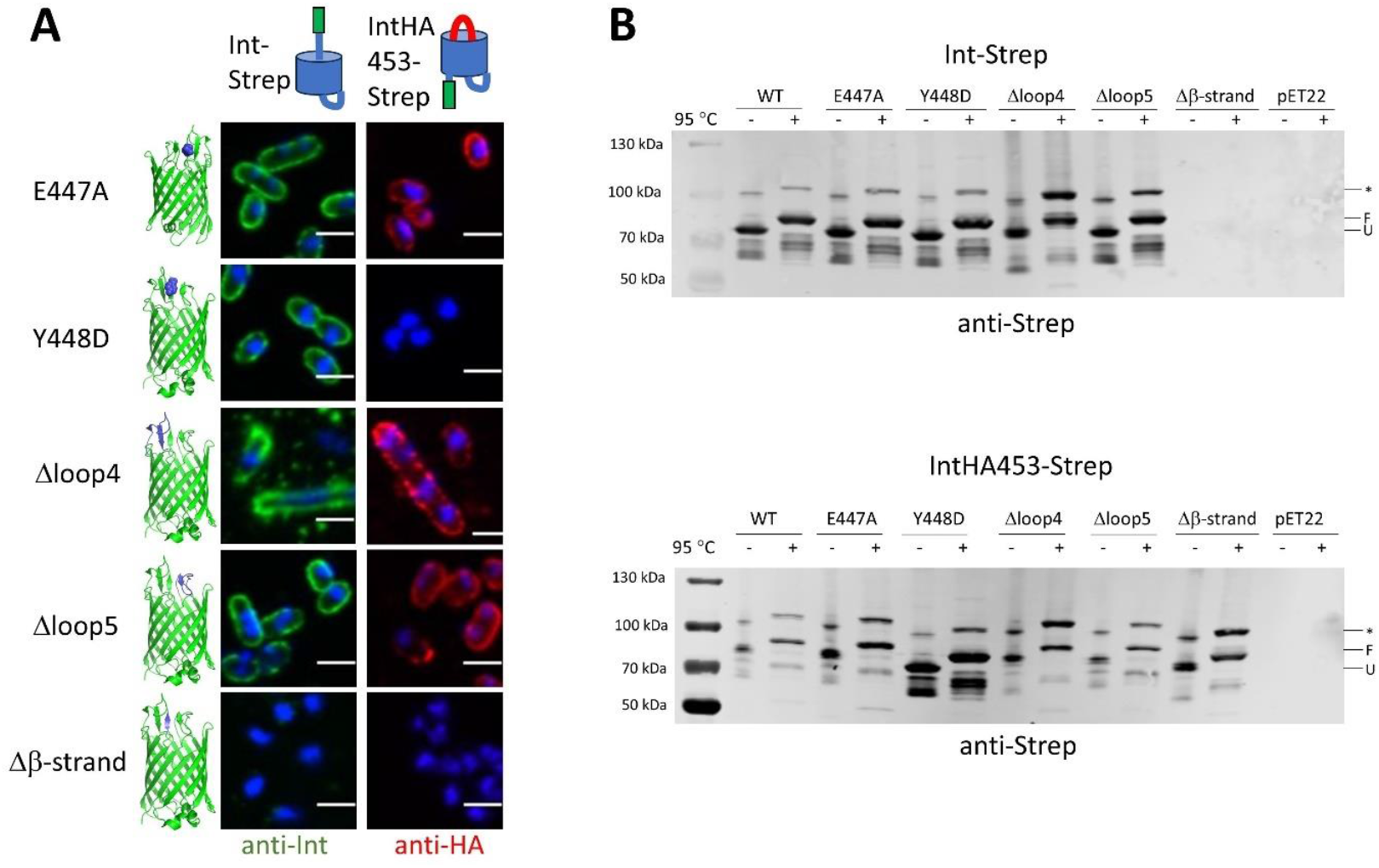
Effect of mutations on extracellular face of intimin. A) Confocal microscopy of intimin mutants in both the Int-Strep (left, probed with an anti-Strep antibody, coloured in green) and IntHA453-Strep (right, probed with an anti-HA antibody, coloured in red) backgrounds. Cells were counterstained with DAPI. Structures on the left highlight the position of the mutation. Schematics at the top show the expected conformation of the proteins. B) Western blots of outer membrane samples with intimin variants using an anti-Strep tag primary antibody. The upper blot is in the Int-Strep background and the lower blot in the IntHA453-Strep background. Samples were split in half and incubated at either room temperature or 95 °C for 10 minutes and run in a 4-12% Novex gel to display the heat shift typical of β-barrel proteins before transferring to a nitrocellulose membrane. pET22 is the empty vector. Molecular weight standards are notated on the left of the gels. F = folded, U= unfolded, * = band of unknown origin.

### Mutants affecting the linker and extracellular β-sheet have reduced surface exposure of the intimin passenger

To quantify the amount of protein exposed at the surface of the intimin mutants demonstrating reduced staining in immunofluorescence microscopy, we performed flow cytometry. Cells expressing Int-Strep or IntHA453-Strep stained well showed good staining with an anti-Strep or anti-HA antibody, respectively (Figure 6). We tested the point mutants Y448D and the mutants affecting larger regions, Δ*α*-helix, Δlinker, Δloop4, Δloop5, and Δβ-strand. In addition, we included two point mutants that had no effect in our microscopy analysis (I382D and R434A) from the periplasmic face of the lumen of the intimin β-barrel, respectively, as controls. The results are largely consistent with what we observed in the fluorescence microscopy: mutants I382D and Δloop5 did not have an appreciably affect (Figure 6A and G) in either the Int-Strep or the IntHA543-Strep backgrounds, whereas deleting the β-strand essentially abolished fluorescence in both (Figure 6H). The Δ*α*-helix variant showed a modest reduction in fluorescence in both cases (Figure 6B), but the Δlinker mutant lost almost all fluorescence in the Int-Strep background, with only a slight reduction in the IntHA453-Strep background (Figure 6D). Interestingly, the R434A mutation resulted in a bimodal distribution in the IntHA453-Strep background, suggesting a slight defect in hairpin formation, but passenger secretion was not impaired (Figure 6C). For the Δloop4 variant, fluorescence was significantly reduced in the Int-Strep background, whereas a bimodal distribution was seen in the IntHA453 background. Quantification of the mean fluorescence intensities (Figure 6I and J) showed that the Δ*α*-helix, Δloop4, and Δβ-strand were all significantly reduced compared with the WT. By contrast, in the IntHA453-Strep set, none of the variants reached significance in a formal statistical test, but the reduction in fluorescence for the Y448D and Δβ-strand variants is clear. The empty vector control was significant in both cases (p < 0.05, not shown in the figure).

**Figure 6.**
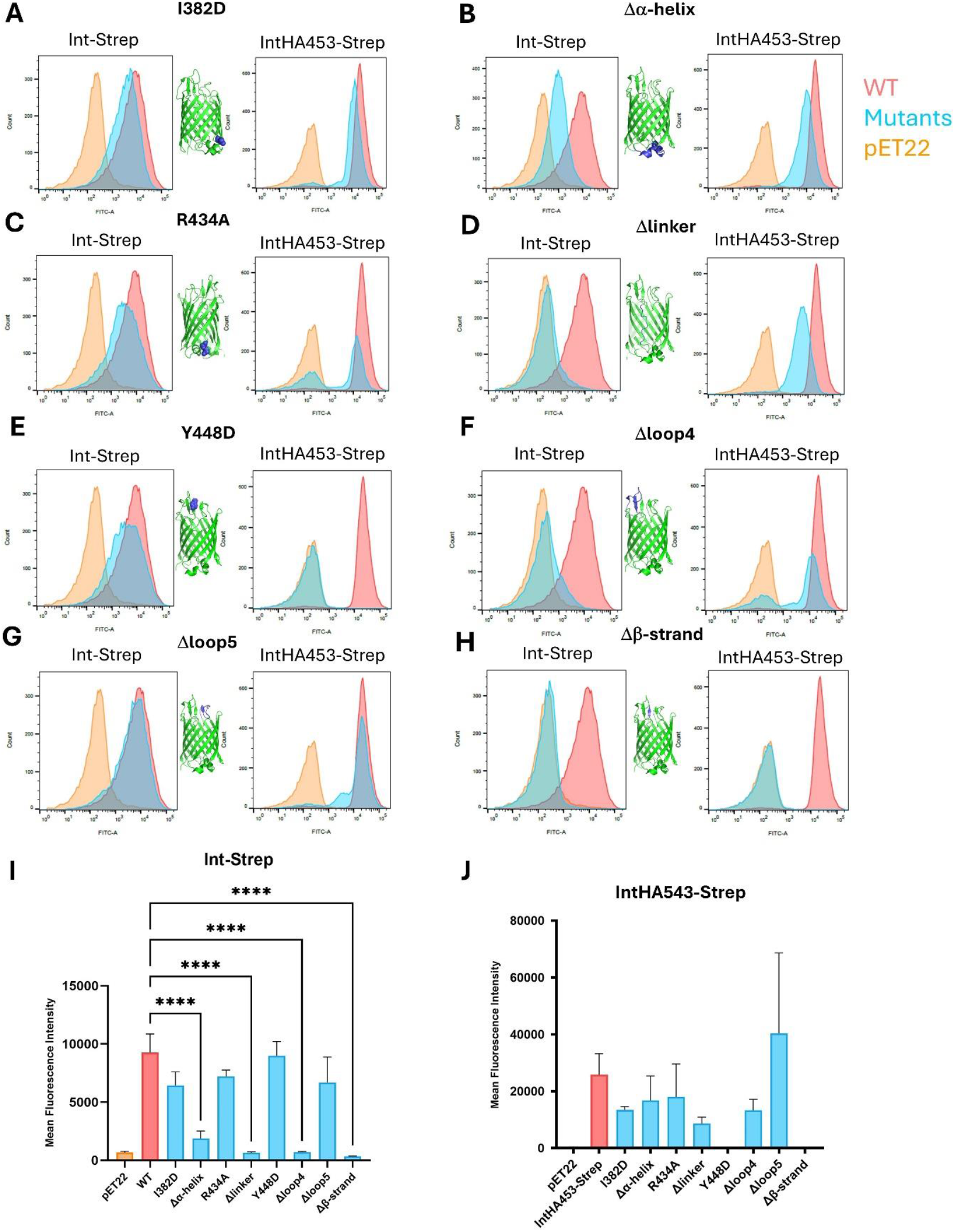
Flow cytometry analysis of selected intimin mutants. The data for the mutants (in both the Int-Strep and IntHA453-Strep backgrounds) are shown by histograms in light blue, the WT is shown in red and the empty vector (pET22) in orange. Bacteria were stained with an anti-Strep primary antibody (for the Int-Strep variants) or an anti-HA antibody (for the IntHA453-Strep variants). Representative histograms (N = 2-3) are shown. A) I348D mutant. B) Δ*α*-helix mutant. C) R434A mutant. D) Δlinker mutant. E) Y448D mutant. F) Δloop4 mutant. G) Δloop5 mutant. H) Δβ-strand mutant. I) Mean fluorescence intensity of cytometry experiments in the Int-Strep background. J) Mean fluorescence intensity of cytometry experiments in the IntHA453-Strep background *** p < 0.001 (one-way ANOVA with Dunnett’s post hoc test; non-significant differences are not indicated).

We reasoned that the reduction of protein levels in the Δβ-strand variants, Δlinker variants and IntHA453-Strep Y448D might be due to loss of the plasmid if expression of these constructs were toxic to the cell. We therefore checked plasmid retention levels by plating dilutions of autoinduced cultures on both LB without selection and LB+ampicillin (Supplementary Figure 7). All overnight cultures expressing intimin displayed plasmid loss, but this was at a similar level and is not enough to explain the complete absence of the protein in the Δβ-strand variants and IntHA453-Strep Y448D. Therefore, any reduction in protein production levels for these variants must be due to either the instability of the resulting proteins or inefficient membrane integration.

### Mutations abolishing secretion also prevent attachment to host cells

To determine whether mutations that affect hairpin formation and secretion had an effect on intimin function, we performed adhesion assays with selected mutants on Tir-expressing HeLa cells (Figure 8). In these assays, bacteria expressing Int-Strep, but not IntHA453-Strep or containing the empty vector pET22, bound to cells primed with Tir. The mutants we tested had displayed some effect on either passenger secretion or hairpin formation in immunofluorescence microscopy (Table 1, Figure 7). Only mutants in the Int-Strep background were tested, as IntHA453-Strep, which does not secrete the C-terminal Tir-binding region to the bacterial surface, is not expected to mediate any adhesion. We also included Int-Strep L437A, which did not have a major effect on either hairpin formation or secretion (Figure 5). Most of the tested mutants bound to Tir-expressing cells at levels comparable to or even higher than the wildtype (Figure 7, Table 1). Reductions were seen for the two mutants where the extracellular loops were removed (Int-Strep Δloop4 and Δloop5), though these results did not reach statistical significance (*p* < 0.15, one-way ANOVA with Dunnett’s multiple comparisons test). However, the Int-Strep Δβ-strand mutant completely failed to mediate adhesion. Adhesion of bacteria expressing the mutants shows that the C-terminal binding domain is correctly folded and presented at the cell surface, though this is reduced for the Δloop4 and Δloop5 mutants, and abolished for the Δβ-strand mutant, which is consistent with the lack of this protein (Figure 5).

**Table 1.**
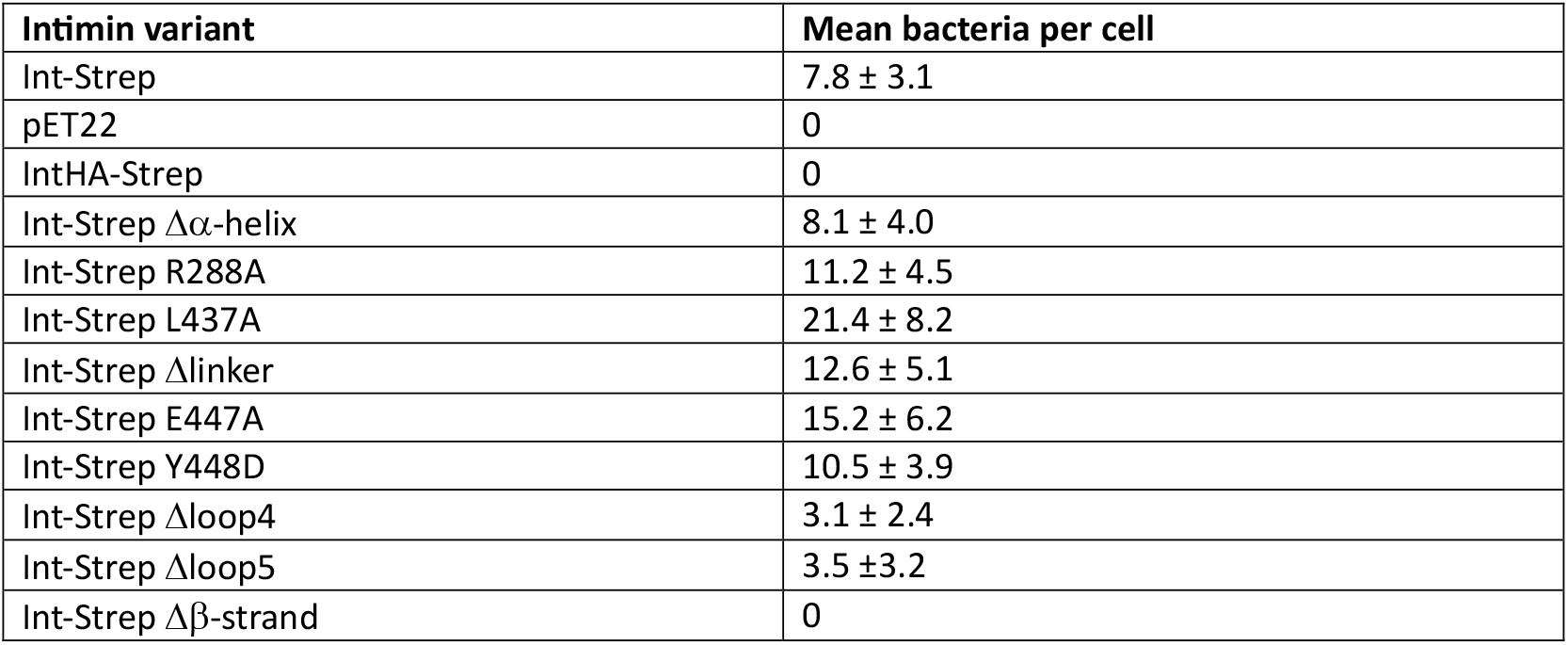
Adhesion of bacteria expressing intimin variants to Tir^+^ HeLa cells. The number of bacteria adhering to 10 infected cells from one adhesion experiment was counted and the mean number of bacteria per cell is shown with standard deviation. All mutants in the table are in the Int-Strep background.

| Intimin variant | Mean bacteria per cell |
| --- | --- |
| Int-Strep | 7.8 ± 3.1 |
| pET22 | 0 |
| IntHA-Strep | 0 |
| Int-Strep $\Delta\alpha$-helix | 8.1 ± 4.0 |
| Int-Strep R288A | 11.2 ± 4.5 |
| Int-Strep L437A | 21.4 ± 8.2 |
| Int-Strep Δlinker | 12.6 ± 5.1 |
| Int-Strep E447A | 15.2 ± 6.2 |
| Int-Strep Y448D | 10.5 ± 3.9 |
| Int-Strep Δloop4 | 3.1 ± 2.4 |
| Int-Strep Δloop5 | 3.5 ± 3.2 |
| Int-Strep $\Delta\beta$-strand | 0 |

**Figure 7.**
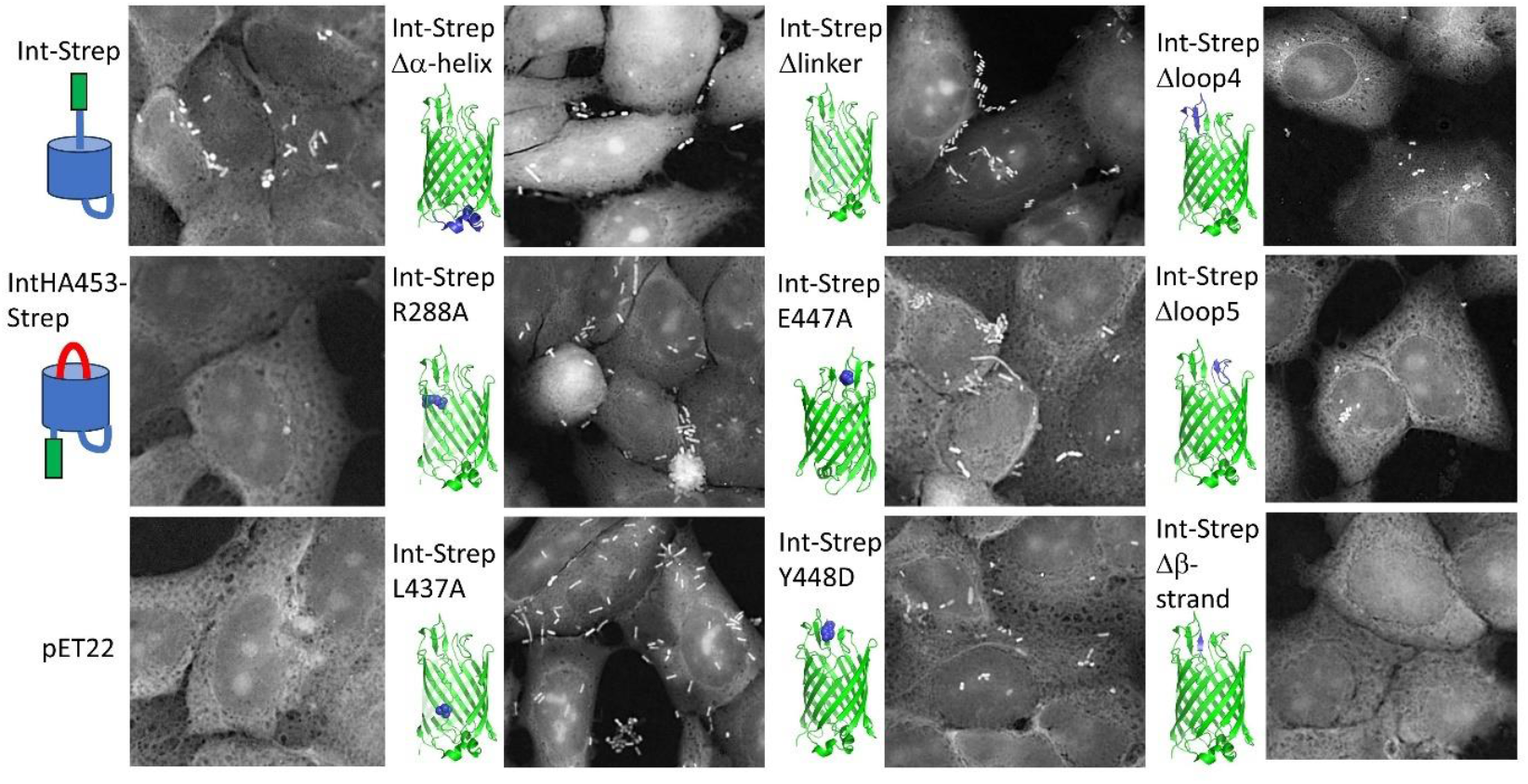
Binding of intimin variants to Tir-displaying HeLa cells. Cells were primed with EPEC lacking intimin, after which BL21ΔF cells producing intimin variants were allowed to adhere. Non-adherent cells were washed off and the cells were stained with fuchsin. The images are shown in negative to highlight the bacteria. Schematics show the expected conformations of control experiments, and the structures highlight the positions of the mutations. pET22 is the empty vector. Quantification of the results is given in Table 1.

## Discussion

The current model for autotransport posits the formation of a hairpin by the linker and the proximal end of the β-barrel, which then leads to vectorial folding and secretion of the passenger [18]. The hairpin mechanism has been long established for classical autotransporters [19-23], and has more recently been shown to be correct for trimeric (type 5c) [24] and inverse (type 5e) [10] autotransporters. In addition, the hairpin mechanism has been suggested for two-partner secretion (type 5b) systems [25], though there is debate about this issue [26,27]. However, how the hairpin is originally formed is mostly unknown.

In trimeric autotransporters, an ‘ASSA’ region was identified within the linker with reduced helical propensity that was hypothesised to be the initiation site for hairpin formation [28]. Introducing proline residues into this region resulted in a partially stalled secretion intermediate [24]. In type 5a secretion systems, a conserved 14-residue sequence in the linker was identified [29]. Mutations in this region revealed that seven of these residues were crucial for passenger secretion. This contrasts with our observations, where individual mutations in the linker or the β-barrel lumen had no effect on hairpin formation or passenger secretion of the inverse autotransporter intimin. Only once the entire linker was changed to a flexible glycine-serine sequence did we see a reduction in surface exposure. However, even then some protein could be secreted and was functional, as shown by adhesion to Tir-expressing cells. This suggests that when the linker is made flexible, a hairpin can still be formed in a subset of cases leading to passenger secretion and stabilisation of the protein. However, the low levels of protein produced in the Int-Strep Δlinker variant indicates that protein where the hairpin is not properly formed is degraded, as is seen for the Δβ-strand variant in the Int-Strep background. Interestingly, in the IntHA453 background the Y448D and Δβ-strand variants were not degraded, suggesting that the double-HA tag somehow protects these proteins from being proteolysed.

The role of the β-barrel in autotransporter biogenesis is also not fully established. Apart from being (part of) a secretion pore, there is evidence for extracellular loop 5 of the classical autotransporter Pet forming a scaffold that acts as a template for passenger folding [30]. In EspP, intrabarrel residues mediate the autoproteolysis of the linker to release the passenger into the extracellular domain [31]. A study investigating the role of conserved residues in a type 5a β-barrel found mutations that had an effect on outer membrane integration and passenger secretion [32]. These were mostly in aromatic residues involved in the so called β-signal needed for BAM recognition [33,34], the aromatic girdle at the hydrophobic membrane interface [35], in a structural mortise-tenon motif [36,37], or suspected to be involved in autoproteolysis. In the trimeric autotransporter YadA and inverse autotransporter intimin, exchanging a conserved glycine in a conserved mortise-tenon motif for larger residues impaired passenger secretion, suggesting a narrowing of the secretion pore [7,15]. Recently, the C-terminal β-strand 12 and its final aromatic residue (part of the β-signal) were shown to be necessary for the biogenesis of the inverse autotransporter YeeJ but replacing β-strand 1 with the sequence of β-strand 3 did not have an effect, showing that the β-barrel can tolerate large changes [38].

Our main finding is that the β-sheet at the extracellular surface of the intimin β-barrel, formed between the linker and extracellular loops 4 and 5, is crucial for intimin stability. When the residues in the β-strand at the C-terminus of the linker were replaced with glycines, the protein was no longer produced. This could either be due to failure at early steps of biogenesis pathway, e.g. interaction with the BAM before β-barrel closure and outer membrane insertion, or at a later stage due to destabilisation of the protein. We prefer the former explanation, as destabilisation would probably lead to reduced protein levels or detectable degradation products, not absence entirely. As no protein could be detected in a DegP-negative strain either, this suggests the problem occurs early in the biogenesis pathway and no protein makes it into the outer membrane.

Again, it is surprising that individual mutations in the β-strand (E447A, Y448D) did not have that great an effect on passenger secretion. The Y448D mutation did, however, lead to loss of hairpin formation – but not to the loss of protein in the outer membrane, suggesting that in this case backsliding of the hairpin into the periplasm could be the explanation for why no HA tag was detected at the bacterial surface. In the IntHA453-Strep background, the mutation Y448D disrupts the hydrophobic cluster at the mouth of the β-barrel, and the introduced aspartate may be excluded from the hydrophobic environment, explaining why the backsliding might occur. In Int-Strep, where the D00 domain is able to fold, folding of the first extracellular Ig-like domain could be enough to prevent backsliding, explaining why the Y448D mutation did not have a large effect in the Int-Strep background. However, even in this scenario the hairpin must form at least transiently to allow for passenger secretion, so any exclusion of Y448D and backsliding must be a relatively slow process compared with the folding of D00, which would prevent any backsliding.

It is also interesting to note that deletion of the individual loops 4 or 5 did not have that great an effect. Loop 4 is the larger of the two loops and removing this reduces the amount of protein displayed on the cell surface. However, for both loop deletion mutants, bacteria were still able to bind to Tir-displaying mammalian cells, albeit at reduced levels, showing that the exported protein is correctly folded. In these variants, the other loop is still in place, and this might be enough to allow formation of a β-sheet between the remaining loop and the β-strand at the top of the linker, allowing for some passenger secretion. We expect that deletion of both loops would lead to abolishing secretion and probably protein stability completely, as seen for the Δβ-strand variant.

There is currently no experimental structure of intimin showing the connection between the β-barrel domain and the first extracellular Ig-like domain, D00. However, in the Alphafold [39,40] model of this region, the connector between the β-strand and the D00 domain is quite long (4 extended residues; see Supplementary Figure 8). If this connector is not projected into the extracellular space by the formation of the small β-sheet the D00 domain might not reach the outside of the β-barrel and be prevented from folding. Why this would lead to protein degradation is unclear, because in the wildtype HA453 variant the D00 cannot fold at the cell surface. The higher stability of the WT IntHA453 variant could be because the β-sheet at the exit of the β-barrel and hairpin are correctly formed. In the absence of the β-sheet the Int-Strep polypeptide might form partly folded structures in the lumen of the (hybrid) β-barrel preventing β-barrel closure and release from the BAM, leading to activation of stress responses (e.g. BepA [41]) that degrade the mis-assembled intimin.

Apart from the extracellular β-sheet, we did not identify interactions within the intimin β-barrel that would lead to the formation of the hairpin. A caveat of our experimental design is that the interactions we targeted are based on the final structure. Therefore, other residues might be involved in transient interactions in the early stages of hairpin formation. However, the robustness of the process, even with large changes such as making the periplasmic *α*-helical turn and linker fully flexible, suggests individual residues are not very important.

In conclusion, we found that the intimin β-barrel domain is largely tolerant of changes and most individual residue exchanges do not affect passenger secretion. Only larger changes, such as replacing the linker traversing the barrel lumen, had noticeable effects on passenger secretion. Importantly, the β-sheet at the extracellular face of the β-barrel appears to be crucial not only for passenger secretion but protein stability overall. This may be because perturbing this region prevents the D00 domain from reaching the outside of the cell to initiate passenger folding and secretion.

## Methods

### Strains and growth conditions

For cloning, *E. coli* TOP10 (Invitrogen) was used. Expression of intimin and its variants was done using *E. coli* BL21ΔF [42], which is equivalent to the BL21(DE3)omp2 strain [43] used in previous studies lacking the abundant porin OmpF for improved outer membrane protein production [7,10,13]. A *degP*-negative version of the BL21ΔF strain (BL21ΔF *degP*) was used to check the effect of DegP [44]. For flow cytometry experiments, BL21Gold(DE3) (Agilent) was employed. For adhesion assays, EPEC E2348/69 Δ*eae* was used [7]. *E. coli* was propagated on lysogeny broth (LB) medium [45] at 37 °C (TOP10) or 30 °C (BL21ΔF). Autoinduction was performed in ZYP-5052 medium [46] for 20 hours at 30 °C. Where necessary, media were supplemented with ampicillin at 100 µg/mL. For adhesion assays, bacteria (EPEC, BL21ΔF) were grown in liver medium [7] at 37 or 30 °C, respectively.

### Cloning and site-directed mutagenesis

To make use of the convenience of autoinduction, we re-cloned intimin and intimin-HA453 from constructs based on the pASK-IBA2 plasmid [7,10] into the pET22b+ (Novagen), with a PelB signal peptide. This was done using Gibson assembly [47]. Briefly, both insert and vector were amplified by PCR (using Q5 polymerase from New England Biolabs; primer sequences are given in Supplementary Table 1) and the resulting products assembled and transformed into TOP10. Transformants were selected on LB+ampicillin at 100 µg/mL and insert-positive clones were screened for using colony PCR. The correctness of the constructs was verified by Sanger sequencing (Eurofins GmbH). Mutations were introduced by a PCR-based site-directed mutagenesis protocol according to [48]. Briefly, primers with overlaps at their 5’ ends incorporating the mutations were designed to amplify the entire plasmid. The template was removed by DpnI digestion (Fermentas), after which the product was transformed into TOP10 cells and transformants were selected for on LB+ampicillin. Mutations were verified by Sanger sequencing (Eurofins GmbH). All primer sequences are given in Supplementary Table 1.

### Outer membrane isolation

Small-scale outer membrane isolation was performed essentially as described in [49]. Briefly, BL21ΔF strains containing intimin expression plasmids were grown overnight in 50 mL autoinduction medium at 30 °C. The following day, an amount of culture equivalent to 40 mL at an optical density at 600 nm (OD_600_) of 1.0 was pelleted and resuspended in 1 mL of lysis buffer (10 mM HEPES pH 7.5, 1 mM MgCl_2_, 1 mM MnCl_2_, 0.1 mg/mL lysozyme and 10 µg/mL of DNase I). The cells were lysed using a bead beater (Precellys Evolution, with 0.1 mm glass beads). Cell debris were pelleted by a 2-minute centrifugation at 15,600 x *g* after which membranes were pelleted by centrifuging for 30 minutes at 20,000 x *g*. The inner membrane was solubilised with 1% *N*-lauroylsarcosine for 30 minutes, after which outer membranes were recovered by centrifuging for 30 minutes at 20,000 x *g*. The membranes were washed once with 10 mM HEPES pH 7.5, after which the outer membranes were resuspended in 40 µL of 10 mM HEPES pH 7.5 and 10 µL of non-reducing 5 x SDS-PAGE loading dye was added. Samples were stored at -20 °C until use.

### Western blots

Outer membrane samples were split in half, and one aliquot was heated for 10 minutes at 95 °C before loading onto an SDS-PAGE gel (Novex gels from Invitrogen). The other half was kept at room temperature before loading. For heat shifts, 4-12% gradient gels were used. The proteins were separated on the gel and then transferred to a nitrocellulose membrane (Whatman Protran) with a Bolt miniblot module (Invitrogen). Once transferred, the membrane was blocked with PBS + 2% skimmed milk powder, either for one hour at room temperature or overnight at 4 °C. The primary antibody (rabbit anti-Strep, Thermo PA5-14454, 1:2000, or rabbit anti-intimin, 1:1000 [7]) was diluted in blocking buffer and incubated with the membrane for 1 hour at room temperature. The membrane was then washed three times with PBS-T (PBS + 0.05% Tween20), after which the secondary antibody (goat anti-rabbit CF770, Biotium) was added diluted 1:10,000 in blocking buffer. After one hour at room temperature, the membrane was washed three times as above and then air dried before imaging with a LI-COR Odyssey CLx imager. For whole cell samples, cells were prepared as in the first steps of outer membrane preparation. After the 2-minute centrifugation, a sample was taken for SDS-PAGE and heated for 10 minutes at 95 °C, after which the procedure was the same as above.

### Immunofluorescence microscopy

For immunofluorescence staining, cultures were auto-induced in 5 ml ZYP-5052 auto-induction medium and grown overnight. About 2 × 10^7^ cells in 1 ml PBS were collected, by measuring OD_600_, on polyethyleneimine-coated 10mm coverslips in a polystyrene 24 well microtitre plate by centrifugation at 4000 x *g* for 5 min. The coverslips were washed with PBS to remove cells that did not stick to the coverslip. The cells were then fixed with 200 μL of 4 % paraformaldehyde in PBS for 30 min. After fixing, the cells were washed and blocked with 1 % Bovine Serum Albumin (BSA) (from VWR chemicals) in PBS at room temperature for 1 hour. The primary antibodies were the rabbit anti-intimin [7] (1:200) and rabbit anti-HA antibody (Santa Cruz Biotechnology, 1:100) in 1 % BSA were added and incubated for 1 hour at room temperature. The coverslips were washed again three times with PBS-T following which CF488A Goat anti-rabbit antibody (1:200, from Biotium) in 1 % BSA was added and incubated in the dark for 2 hours at room temperature. The coverslips were then mounted on a glass slide with 10 μl of Biotium’s EverBrite™ Mounting Medium with DAPI and sealed around the perimeter with Biotium’s CoverGripTM Coverslip Sealant. The samples were imaged using an UPlanFLN 100x/1.3 oil immersion on an Olympus Fluoview FV1000 Inverted Confocal Microscope. The blue channel (DAPI) was imaged first and then the green channel (CF488A).

### Flow cytometry

For flow cytometry experiments, *E. coli* BL21Gold(DE3) cells with intimin plasmids were grown overnight in 5 mL autoinduction medium supplemented with ampicillin at 37 °C. The cultures were washed with PBS, after which the bacteria were resuspended in 5 mL PBS and the OD_600_ was measured. 2 × 10^8^ bacteria were pelleted and resuspended in 600 µL of PBS, after which 600 µL of 8% formaldehyde was added and the suspensions incubated for at least 30 minutes at room temperature. The fixed bacteria were then washed with PBS, after which the cells were resuspended in 1 mL PBS with 1% BSA and blocked for one hour at room temperature. The primary antibody (rabbit anti-Strep (Abcam), 1:200 or rabbit anti-HA (Cell Signalling Technology), 1:500) was then added and the cells incubated with rotation overnight at 4 °C. The following day, the cells were washed once with PBS, after which the secondary antibody (anti-rabbit CY2 (Abcam), 1:200 in PBS + 1% BSA) was added. The cells were incubated for 1 hour with rotation in the dark at room temperature. The bacteria were washed once with PBS and then resuspended in 300 µL PBS + 1% BSA. Pre-blocked Falcon 5 mL round-bottom tubes were used to inject the bacteria into the flow cytometer FSC-A, SSC-A, and FITC-A signals were recorded in arbitrary units (a.u.) using a BD LSRFortessa™ and BD FACSDiva™. Detector voltages were set to 489 V (FSC), 225 V (SSC), and 560 V (FITC). A minimum of 10.000 events per sample were collected within a defined gate based on FSC-A vs SSC-A. The results were analysed using Flowjo^™^ (by FlowJo, LLC, a BD company).

### Plasmid stability assays

To assay for potential toxicity of intimin constructs and consequent loss of plasmid, we grew BL21ΔF with intimin constructs or the empty vector pET22b+ in autoinduction medium overnight at 30 °C. The following day, the OD_600_ of the cultures was measured and cultures were adjusted to the same OD_600_. From this, bacteria were serially diluted in sterile PBS and 5 µL of dilutions (10^0^-10^−6^) were spotted onto LB or LB + ampicillin 100 µg/mL. Plates were incubated overnight at 30 °C and imaged.

### Adhesion assays

Adhesion of intimin-expressing bacteria to HeLa cells was assessed using a prime-challenge assay, essentially as described in [7]. Briefly, HeLa cells were pre-infected with EPEC E2348/69 Δ*eae* at an multiplicity of infection (MOI) of 100 to inject them with Tir. The non-adherent cells were washed off the cells, and any remaining bacteria were killed with gentamicin. After further washes, BL21ΔF containing intimin variants and induced with 0.5 mM isopropylthiogalactoside for 4 hours were added to the cells at an MOI of 100 for 1 hour. The cells were then washed with PBS until the negative control strain showed no adhesion of bacteria to cells in light microscopy. The samples were then fixed with 4% paraformaldehyde (overnight at 4 °C or 1 hour at room temperature), washed and stained with Fuchsin for approximately 30 seconds. The coverslips were washed and air dried and then mounted onto glass slides with the mounting medium Roti® Histokitt. The cells were viewed under an Olympus light microscope at 60 x magnification with oil immersion and imaged using Cell B software.

## Data availability statement

The microscopy images generated in this study are available through FigShare (10.6084/m9.figshare.28945613). All other data are included in this paper.

## Acknowledgements

We thank Dirk Linke (University of Oslo) for support and discussions. Victoria Wootton (Nottingham Trent University) is acknowledged for assistance with western blots. The microscopy presented in this study was performed at the Oslo NorMIC Imaging Platform at the Department of Biosciences, University of Oslo. Members of the Antimicrobial Resistance, Omics and Microbiota group at Nottingham Trent University are thanked for discussions and collegiality.

