## Supplementary material for "Formation of the β-sheet at the extracellular opening of the intimin β-barrel domain is necessary for stability and efficient passenger secretion"

### Supplementary Material for Sarma et al.

**Table 1.** Primers used in this study.

| Primers | Sequences (5'→3') | Comment |
|---|---|---|
| pET22rec Fwd<br>pET22rec Rev | TGAGATCCGGCTGCTAACAAAG<br>GGCCATCGCCGGCTGG | For amplifying pET22 for cloning |
| pelB-Int Fwd<br>pET22-Strep Rev | GCCCAGCCGGCGATGGCCAATGGTGAAAATTATTTTAAATTGGGTTC<br>GTTAGCAGCCGGATCTCATTATTTTTCGAACTGCGGGTGG | For amplifying intimin (with C-terminal StrepII tag for cloning) |
| IntD298A Fwd<br>IntD298A Rev | AATACTGGCGAGCTTATTTCAAAAGTAGTGTTAACGGCTA<br>AGCTCGCCAGTATTCGCCACCAATACCTAAACGG | Mutagenesis primers to introduce the D298A mutation |
| IntI382D Fwd<br>IntI382D Rev | AACTATACTCCGGATCCTCTGGTGACGATGGGG<br>ATCCGGAGTATAGTTTACACCAACGGTCGCCGC | Mutagenesis primers to introduce the I382D mutation |
| IntP421A Fwd<br>IntP421A Rev | CAGCAAATTGAGGCTCAATATGTTAACGAGTTAAGAACATTAT<br>AGCCTCAATTTGCTGGGACCACGGTTTATCAAACTGAT | Mutagenesis primers to introduce the P421A mutation |
| IntL430A Fwd<br>IntL430A Rev | GAGTTAAGAACAGCTTCAGGCAGCCGTTACGATCT<br>AGCTGTTCTTAACTCGTTAACATATTGTGGCTCAATTTGC | Mutagenesis primers to introduce the L430A mutation |
| IntΔα-helix Fwd<br>IntΔα-helix Rev | TTTTCTGGTTCTGGTTCAGGCAGCCGTTACGATCT<br>ACCAGAACCAGAAAACTGATAACGGAACTGCATTGAGT | Mutagenesis primers to replace residues 412-430 with SGSG |
| IntF276A Fwd<br>IntF276A Rev | GCTATAACGTCGCCATTGATCAGGATTTTTCTGGTGATAAT<br>GGCGACGTTATAGCCCAACATATTTTCAGGAAGGAAAAAA | Mutagenesis primers to introduce the F276A mutation |
| IntR288A Fwd<br>IntR288A Rev | GGTGATAATACCGCGTTAGGTATTGGTGGCGAATACTG<br>CGC GGT ATT ATC ACC AGA AAA ATC CTG ATC AAT GAA GAC G | Mutagenesis primers to introduce the R288A mutation |
| IntR434A Fwd<br>IntR434A Rev | TCAGGCAGCGCTTACGATCTGGTTCAGCGTAATAA<br>AGCGCTGCCTGATAATGTTCTTAACTCGTTAACATATTGT | Mutagenesis primers to introduce the R434A mutation |
| IntL437A Fwd<br>IntL437A Rev | GCCGTTACGATGCTGTTCAGCGTAATAACAATATTATTCTG<br>AGCATCGTAACGGCTGCCTGATAATGTTCTTAACTCGTT | Mutagenesis primers to introduce the L437A mutation |
| IntQ439A Fwd<br>IntQ439A Rev | ACGATCTGGTTGCTCGTAATAACAATATTATTCTGGAGTAC<br>AGCAACCAGATCGTAACGGCTGCCTGATAATGTTCTT | Mutagenesis primers to introduce the Q439A mutation |
| IntΔlinker Fwd<br>IntΔlinker Rev | AGTGGGTCTGGCTCCGGTAGCATTCTGGAGTACAAAAAGCAGGATA<br>GCCAGACCCACTACCGCTGCCACGGCTGCCTGATAATGTTCTT | Mutagenesis primers to replace residues 435-444 with GSGSGSGSGS |
| IntE447A Fwd<br>IntE447A Rev | ACAATATTATTCTGGCGTACAAAAAGCAGGATATTCTTTCTC<br>CGCCAGAATAATATTGTTATTACGCTGAACCAGATCGTAA | Mutagenesis primers to introduce the E447A mutation |
| IntY448D Fwd<br>IntY448D Rev | ATTATTCTGGAGGACAAAAAGCAGGATATTCTTTCTCTGA<br>GTCCTCCAGAATAATATTGTTATTACGCTGAACCAGATC | Mutagenesis primers to introduce the Y448D mutation |
| IntΔloop4 Fwd<br>IntΔloop4 Rev | GGTGGGGGTGGGAATGGCTTCGATATCCGTTTTAAT<br>CCCACCCCCACCGAAATAGCCGTTAACACTACT TTT G | Mutagenesis primers to replace residues 309-327 with GGGG |
| IntΔloop5 Fwd<br>IntΔloop5 Rev | GGTGGACCTGGTGCGGCGACCGTTGGTGTA<br>ACCAGGTCCACCTCCACCATACTGCTCATACATCA | Mutagenesis primers to replace residues 354-369 with GGGG |
| IntΔβ-strand Fwd<br>IntΔβ-strand Rev | GGTGGGGGTGGGAAGCAGGATATTCTTTCTCTGAATATT<br>CCCACCCCCACCAATAATATTGTTATTACGCTGAACCAG | Mutagenesis primers to replace residues 446-449 with GGGG |

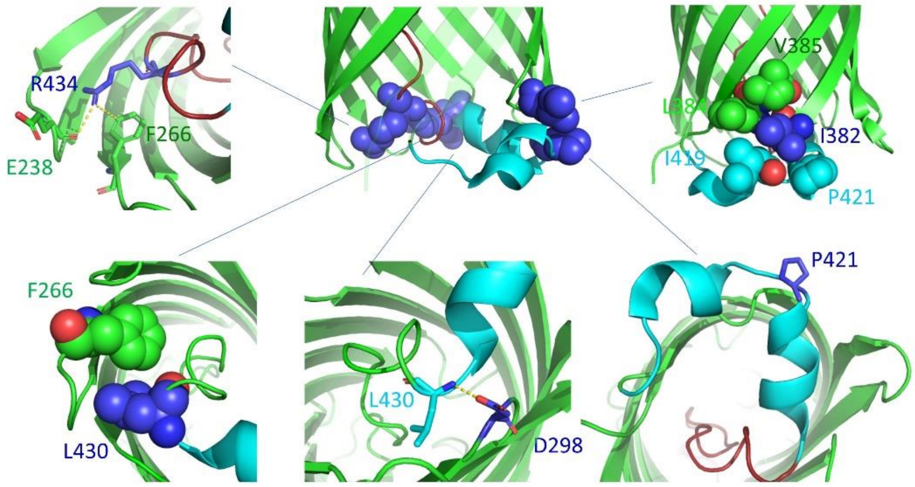


| Mutation | Interactions |
|---|---|
| D298A | L430 |
| I382D | L384, V385, I419, P421 |
| P421A | Potential kink induced |
| L430A | F266 |
| R434A | E238, F266 |

**Supplementary Figure 1.** Residues and interactions targeted at the periplasmic side of the intimin β-barrel domain. Mutated residues are shown in blue, in either space-filling or stick representation; interacting residues are coloured according to the colour scheme in Figure 1 in the main paper, and hydrogen or ionic bonds in dashed yellow lines. D298 interacts with the amide nitrogen of L430, potentially stabilising the N-terminus of the linker. I382 forms the core a hydrophobic pocket involving L384, V385, I419 and P421. P421 is situated in a turn between two short α-helices and may be important in introducing a kink at this position. L430 has a hydrophobic interaction with F266, which may be important in positioning the N-terminus of the linker. R434 in the linker has an ionic interaction with E238 in a periplasmic turn and a cation-pi interaction with F266 in the lumen wall. These interactions are summarised in the table on the right. The Δα-helix mutation replaces the entire helical turn (in cyan in the middle) with a glycine-serine stretch of the same length. The figures were prepared using PyMOL (Schroedinger).

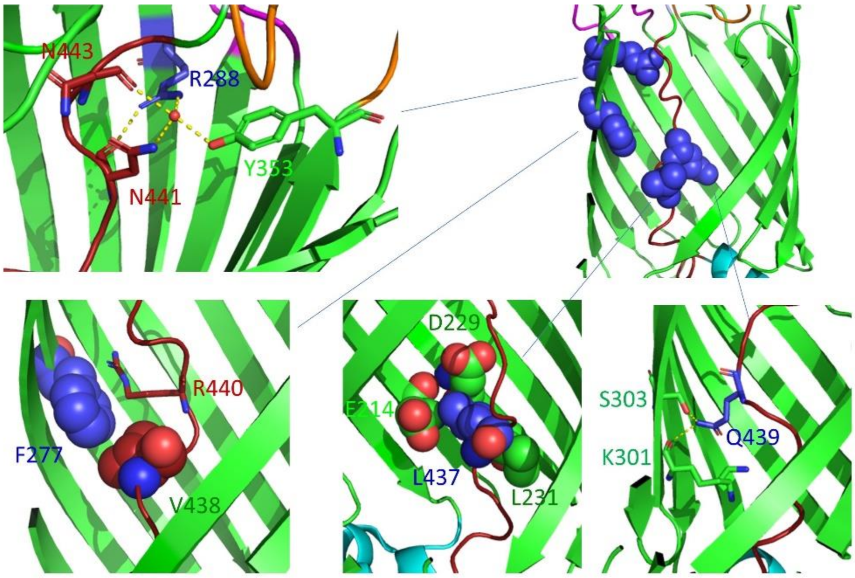


| Mutation | Interactions |
| --- | --- |
| D277A | V438, R440 |
| R288A | Y353, N441, N443 |
| L437 | E214, D229, L231 |
| LQ439 | K301, S303 |

**Supplementary Figure 2.** Residues and interactions targeted in the linker region of intimin. Mutated residues are shown in blue, in either space-filling or stick representation; interacting residues are shown according to the colour scheme in Figure 1 in the main paper, and hydrogen or ionic bonds in dashed yellow lines. F277, located on the lumen wall, has a hydrophobic interaction with V438 as well as cation-pi interaction with R440. R288, also on the lumen wall, interacts through with several residues including N441 and N443 in linker either directly or through a water bridge. L437 in the linker packs against L231, E214 and D229, located in the lumen wall. Q439 in the linker forms hydrogen bonds with K301 and S303 on the lumen wall. These interactions are summarised in the table on the right. In addition, the entire linker (434-444) was replaced by a GS sequence of the same length. The figures were prepared in PyMOL (Schroedinger).

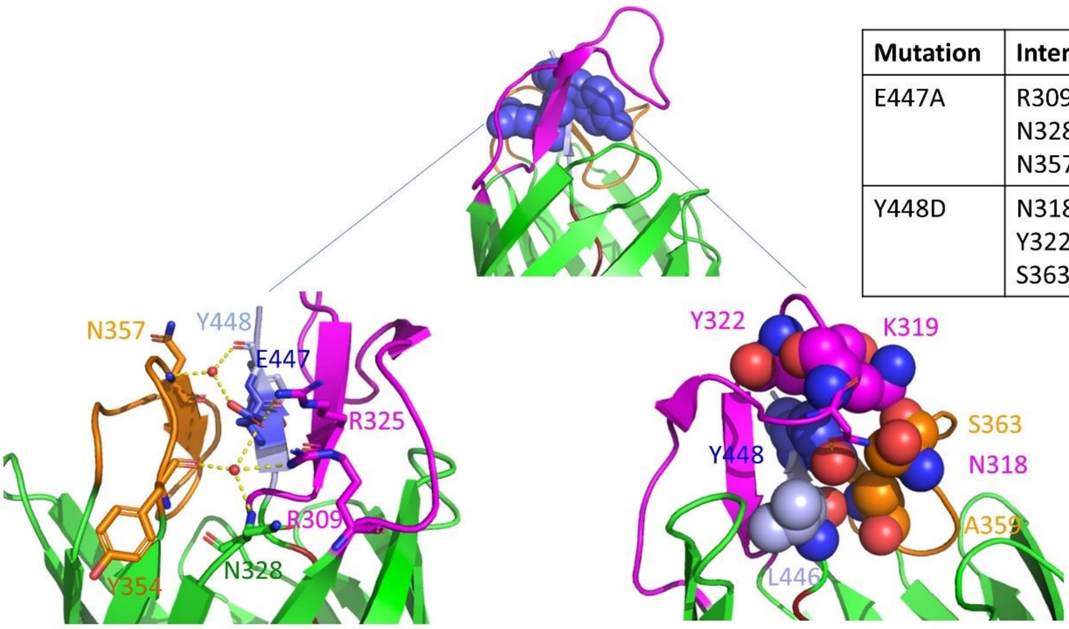


| Mutation | Interactions |
|---|---|
| E447A | R309, R325, N328, Y354, N357, Y448 |
| Y448D | N318, K319, Y322, A359, S363, L446 |

**Supplementary Figure 3**. Residues and interactions targeted at the extracellular region of intimin. Mutated residues are shown in blue, in either space-filling or stick representation; interacting residues are shown according to the colour scheme in Figure 1 in the main paper, and hydrogen or ionic bonds in dashed yellow lines. E447, in the β-strand at the C-terminal end of the linker, is at the centre of an interaction network between several residues in the neighbouring loops. Y448 is at the core of a hydrophobic cluster involving residues from loop 4 (N318, K319, Y322) loop 5 (A359, S363) and the linker (L446). These interactions are summarised in the table on the right. In addition, both loops 4 and 5 were replaced by a four-glycine stretch (Δloop4 and Δloop5, respectively), as was the β-strand itself (Δβ-strand). The figures were prepared in PyMOL (Schroedinger).

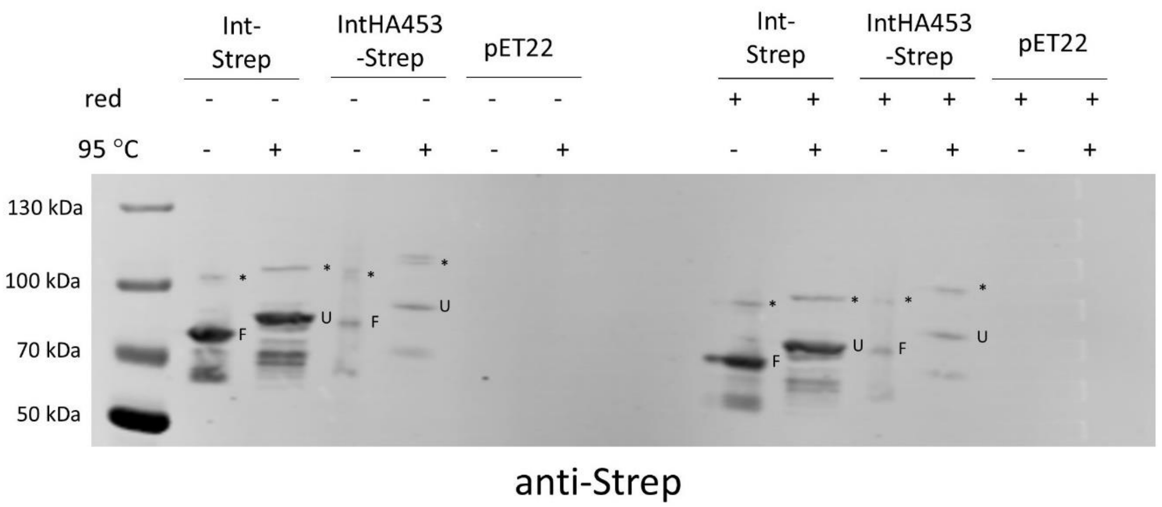


**Supplementary Figure 4**. The additional intimin band is not due to incorrect disulphide bond formation. Outer membrane samples with Int-Strep or IntHA453-Strep were split into two, and to one sample reducing agent (red, Thermo Fisher NuPAGE sample reducing agent) was added. The samples were then split again, and one half was incubated at room temperature while the other was heated for 10 minutes at 95 °C. The samples were then run in a 4-12% Novex gel before transferring to a nitrocellulose membrane and probing with an anti-Strep antibody. pET22 is the empty vector. Molecular weight standards are notated on the left. F = folded, U= unfolded, * = band of unknown origin.

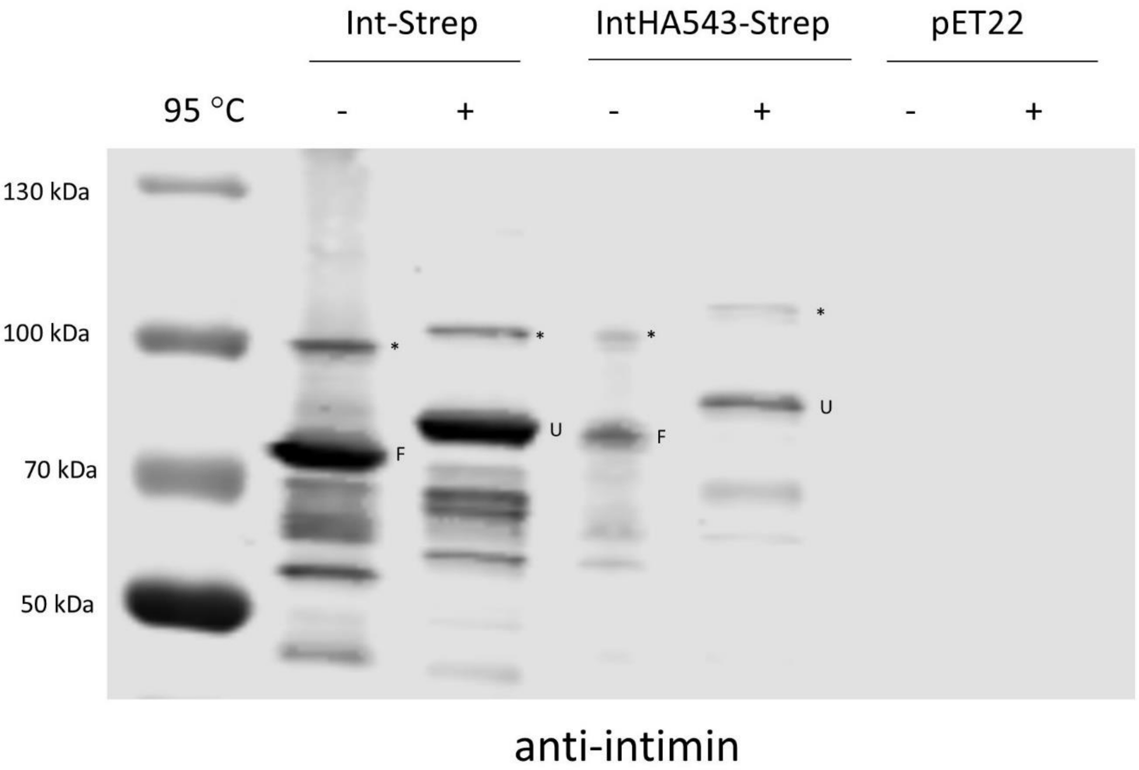


**Supplementary Figure 5**. Western blot of Int-Strep and IntHA453-Strep with an anti-intimin antibody recognising the C-terminus of intimin. Outer membrane samples with Int-Strep or IntHA453-Strep were split into two and one half was incubated at room temperature while the other was heated for 10 minutes at 95 °C. The samples were then run in a 4-12% Novex gel before transferring to a nitrocellulose membrane and probing with an anti-Intimin antibody [7]. pET22 is the empty vector. Molecular weight standards are notated on the left. F = folded, U= unfolded, * = band of unknown origin.

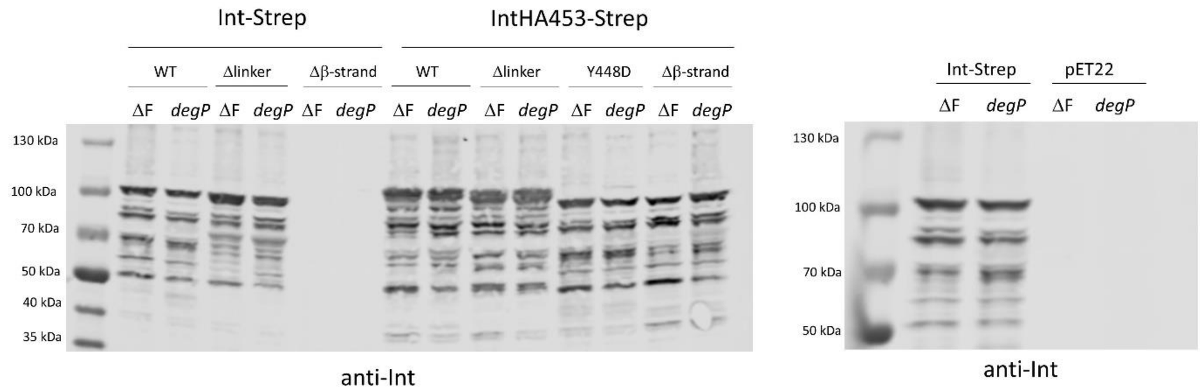


**Supplementary Figure 6.** Loss of protein in the Int-Strep Δβ-strand variant is not due to degradation by DegP. Whole-cell samples from autoinduced cultures of either BL21ΔF (ΔF) or BL21ΔF Δ*degP* (*degP*) expressing intimin variants were separated on a 4-12% gel and transferred to a nitrocellulose membrane before being probed with an anti-intimin antibody [7]. The blot on the left shows intimin variants in either the Int-Strep of the IntHA453-Strep backgrounds. The gel on the right is a control gel where pET22 is the empty vector. Molecular weight standards are notated on the left of the gels.

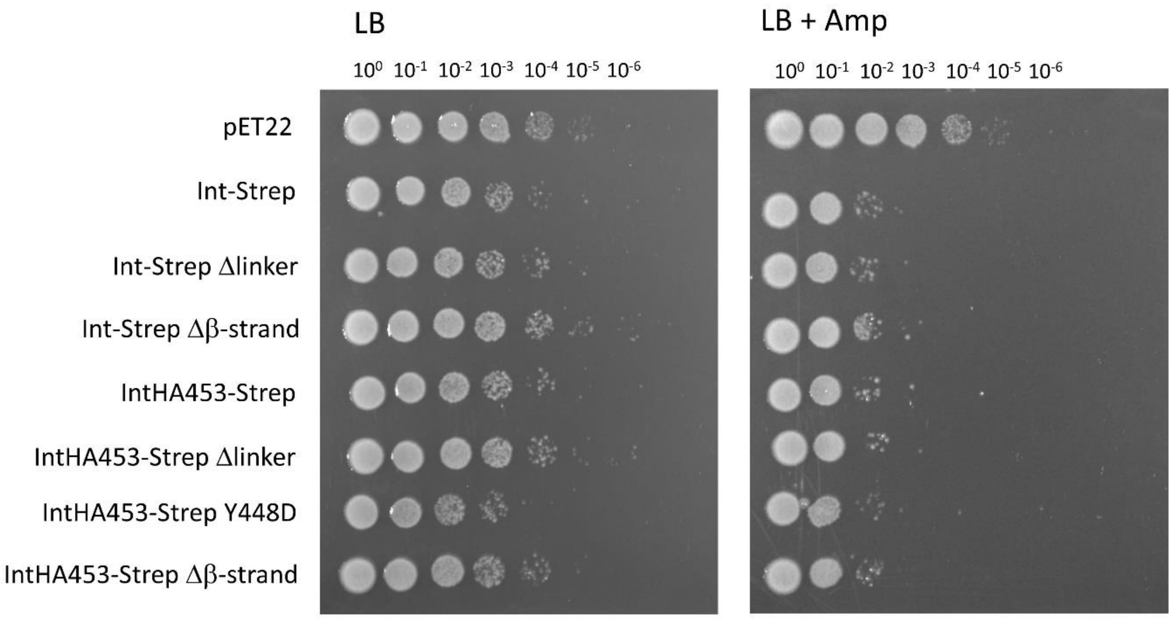


**Supplementary Figure 7**. Plasmid retention in cultures expressing intimin variants. Bacteria were cultured in autoinduction medium overnight and adjusted to the same optical density the following day. Serial dilutions were made and plated onto LB with no selection (left) or LB with ampicillin (right); bacteria lacking the plasmid will not grow on the ampicillin plate. Intimin variants show a similar level of plasmid loss in overnight cultures.

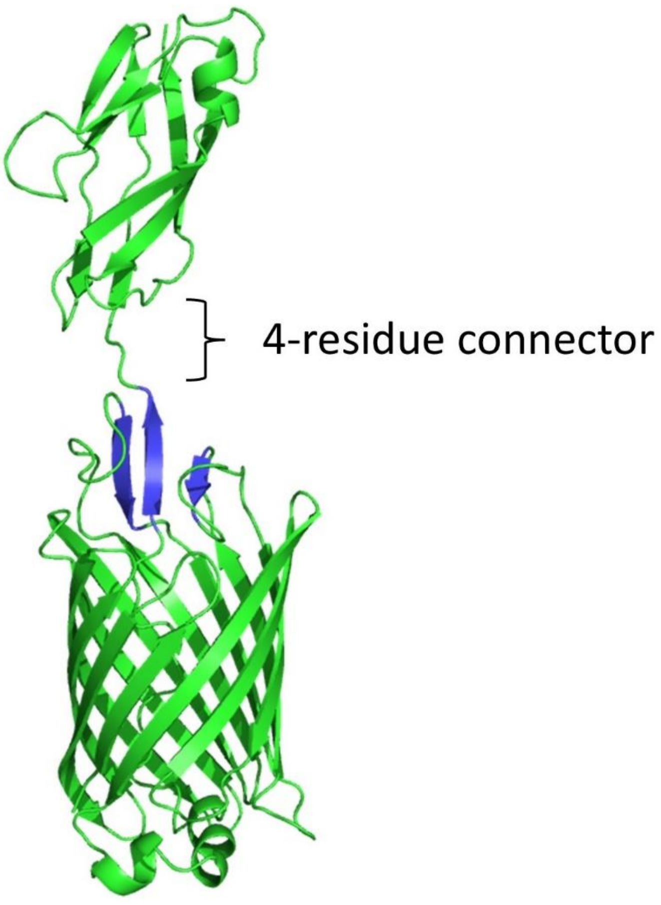


**Supplementary Figure 8**. Alphafold model of the intimin β-barrel domain and D00 domain. The model shows a 4-residue extended connector element between the β-sheet at the extracellular face of the β-barrel (in blue) and the Ig-like D00 domain.
